# Fastq-pair: efficient synchronization of paired-end fastq files

**DOI:** 10.1101/552885

**Authors:** John A. Edwards, Robert A. Edwards

## Abstract

Paired end DNA sequencing provides additional information about the sequence data that is used in sequence assembly, mapping, and other downstream bioinformatics analysis. Paired end reads are usually provided as two fastq-format files, with each file representing one end of the read. Many commonly used downstream tools require that the sequence reads appear in each file in the same order, and reads that do not have a pair in the corresponding file are placed in a separate file of singletons. Although most sequencing instruments capable of generating paired end reads produce files where each read has a corresponding mate, many downstream bioinformatics manipulations break the one-to-one correspondence between reads, and paired-end sequence files loose synchronicity, and contain either unordered sequences or sequences in one or other file without a mate. Trivial solutions to this problem require reading one or both of the DNA sequence files into memory but quickly become limited by computational resources for moderate to large sized sequence files that are common nowadays. Here, we introduce a fast and memory efficient solution, written in C for portability, that synchronizes paired-end fastq files for subsequent analysis and places unmatched reads into singleton files.

Fastq-pair is freely available from https://github.com/linsalrob/fastq-pair and is released under the MIT license.

## fastq-pair

Fastq-format files have become the dominant file format for sharing DNA sequences as they conveniently contain both the sequence and quality information in a single file (1). In addition, paired end sequencing has come to dominate sequencing approaches because of the additional information gained from knowing the distance between the pairs and potentially the longer sequences that can be acquired from by joining the overlapping pairs. For example, the sequence read archive contains twice as many paired end libraries as single read libraries (As of Feb 1st 2018 there were 2,233,015 paired end libraries and 1,110,884 single read libraries in the SRA).

Many downstream tools that are used to join paired end read data (e.g. pear (2)), assemble DNA sequences (e.g. spades or meta-spades (3)), map reads to reference sequences (e.g. bowtie2 (4)), require that the paired sequences be synchronized. In particular these tools require that the two files that represent a single paired end sequencing run (i) have the same number of reads in each file and (ii) that the left and right (or forward and reverse, depending on your preferred terminology) sequences appear in the same order in each file. In contrast, several upstream applications do not provide paired-end sequences in synchronized files. For example, fastq-dump that is widely used to retrieve sequences from the sequence read archive (SRA) does not automatically synchronize the order of sequences in files and several trimming programs result in unordered paired end reads, although an undocumented option (--split-3) maybe appended to the fastq-dump command to request the sequences be split in order (5).

As sequencing technology improves, fastq files are increasing in size, and files of many tens of gigabytes are not uncommon. This provides a challenge for resynchronizing paired end reads so that their reads follow the same order. We developed fastq-pair specifically to handle large fastq files in a memory and time efficient manner.

We wrote fastq-pair to accept two fastq format files as input, and to generate four fastq files as the output. As per the standard (1), the sequence identifier in a fastq file consists of all of the non-whitespace characters on the identifier lines denoted by the leading @ symbol. Forward and reverse reads are typically identified by a trailing /f or /r on the sequence identifier, and left and right are typically identified by a trailing /1 and /2 on the sequence identifier. As most other software does, we currently presume that all fastq files do not wrap either the sequences or the quality information, and thus each sequence is represented by exactly four lines (the identifier line beginning with the @ symbol, the DNA sequence, the spacer line that begins with a + and indicates the end of the DNA sequence, and the quality score line) (1).

Fastq-pair is instantiated by providing the file names for two files. The algorithm creates a hash of objects that contain the sequence identifier and the position in the file for all the identifiers in the first file. This significantly reduces the memory requirements for fastq-pair compared to trivial solutions that store sequence identifier, sequence, and quality scores. The file pointer to the first file remains accessible while the second file is read, and for each sequence in the second file, if the identifier is present in the hash the two sequences and their quality scores are written to the appropriate files and a flag is set to mark that the sequence has been seen. If the identifier is not present in the hash the sequence is written to the appropriate singletons file. After iterating through the second file, any sequences that were not seen are written to the appropriate singletons file.

The algorithm has linear complexity for both memory consumption and execution time (Fig. 1). The size of the hash table may be altered by the user, and a larger table, while consuming more memory and taking slightly longer to instantiate will reduce look-up time to determine whether an identifier is in the hash. The default table size (100,003) is designed to enhance even distribution of elements through the table, and provides linear complexity even for large sequence files.

**Fig. 1.**
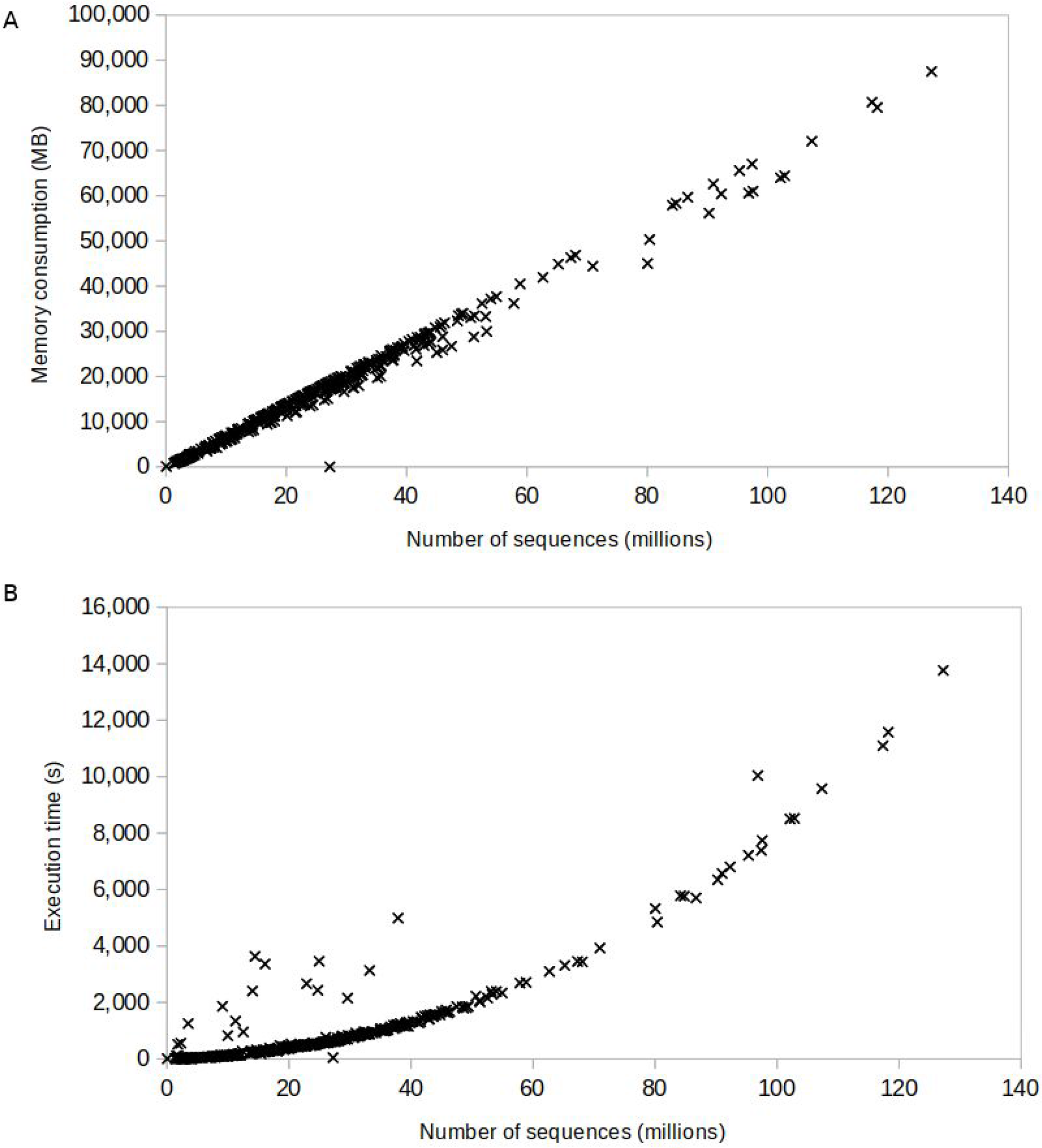
Empirical complexity analysis of the execution of fastq-pair. 422 fastq files from the sequence read archive, with sizes ranging from 66,854 to 127,235,892 sequences were processed with fastq-pair and the time and memory complexity measured using the unix application *time*. Time is the sum of user and system time in seconds, and memory is the maximum resident memory in mb.

The output from fastq-pair is four separate files. The synchronized files are named as the input with *.paired.fq* appended to the file name. The singleton files are named as the input with *.single.fq* appended to the file name. It was a deliberate decision to maintain the two singleton files separately rather than create a single file as (i) this allows a rapid method of determining how many singleton reads there were from each file, and (ii) it lends itself to parallel processing of many files simultaneously as the file names are unique.

There is one caveat: fastq-pair relies heavily on random access to the file stream, and therefore cannot process compressed files. At least one file would have to be decompressed into either memory or a temporary file which defeats the purpose of fastq-pair being memory efficient.

### Conclusion

fastq-pair provides a rapid, memory and time efficient approach for ensuring fastq-files contain the reads in the same order as required for many downstream processing steps.

## Acknowledgements

We thank Vito Adrian Cantu and Katelyn McNair for comments on the manuscript. This work was supported by National Science Foundation grant number MCB-1441985 to RAE.

